# Age-specific modulation of gut-brain axis metabolites by galacto-oligosaccharides and nutrient blends in early childhood

**DOI:** 10.64898/2026.01.29.702544

**Authors:** Laurent Ferrier, Shaillay Kumar Dogra, Lam Dai Vu, Alexandros K. Kanellopoulos, Jonas Poppe, Laurence Biehl, Aurélien Baudot, Pieter Van den Abbeele

## Abstract

**Objectives:** Gut microbiome-derived metabolites, including short-chain fatty acids (SCFA) and tryptophan derivatives, are key mediators of the gut-brain axis. We examined how early-life nutritional interventions influence these metabolites during critical neurodevelopmental periods.

**Methods:** Using a standardized ex vivo fermentation system, we assessed the effects of galacto-oligosaccharides (GOS), nutrient blends (vitamins, minerals, amino acids), and their combinations on the gut microbiome of infants (2–4 months, n=6) and young children (2–3 years, n=6).

**Results:** Baseline microbiome composition differed by age: infants showed low α-diversity and high interpersonal variability, while young children exhibited more adult-like profiles. Nutrient blends increased propionate/butyrate ratios and branched-chain fatty acids (BCFA) in young children, alongside B-vitamin and amino acid-derived metabolites, including neuroactive compounds (indole-3-carboxaldehyde, imidazoleacetic acid, pipecolinic acid). Combining nutrient blends with GOS produced synergistic effects on propionate (infants) and 2-hydroxyisocaproic acid (HICA, both groups). GOS-containing treatments strongly promoted Bifidobacteriaceae, driving production of acetate, HICA, N-acetylated amino acids, aromatic lactic acids, and acetylagmatine; in young children, also butyrate and γ-aminobutyric acid (GABA).

**Discussion:** GOS alone and combined with nutrient blends modulated microbiome-derived metabolites linked to the gut-brain axis. Synergistic effects on GABA, acetylagmatine, and HICA suggest roles in neurotransmission, neuroprotection, and immune–brain signaling. Despite shared bifidogenic effects, age-specific differences indicate developmental stage influences intervention outcomes. Further studies should explore neurodevelopmental benefits of these combinations and metabolites.

## Introduction

The gut-brain axis (GBA) refers to the connection between the gastrointestinal tract and the central nervous system, through neural, endocrine, immune and humoral pathways^1^. It is impacted by the gut microbiome in a bidirectional manner, i.e., the gut microbiome signalling to the brain and *vice versa*. Mechanistically, brain function is impacted by microbial metabolites like short-chain fatty acids (SCFA, e.g. acetate, propionate and butyrate)^2^ and tryptophan-derived metabolites (e.g. indoles)^3^ along with microorganism-associated molecular patterns (MAMP, e.g. nucleotides, lipids, carbohydrates and peptides)^4^. Conversely, the brain can influence gut physiology and function by regulating immune responses and controlling the release of endocrine hormones and neurotransmitters^1^. These signalling molecules travel through systemic circulation, affecting gut motility, immune activity, and microbial composition^5^.

The first few years are crucial for neurodevelopment^6^ and are a period during which the gut microbiome also undergoes drastic alterations, amongst others driven by mode of delivery, type of feeding (breast or formula) and use of antibiotics^7^. While the infant microbiome has a low α-diversity and mainly consists of *Bifidobacteriaceae, Bacteroidaceae* and *Enterobacteriaceae*, the microbiota of young children is highly diverse and dominated by adult-like taxa such as *Bacteroidaceae*, *Lachnospiraceae* and *Ruminococcaceae*^8^. Deviations in gut microbiome composition in early live have been associated with autism spectrum disorder^9^, emphasizing a window of opportunity for nutritional interventions in early life to prevent neurological and mental diseases by impacting the gut microbiome.

Probiotics (live microorganisms)^10^ are a first type of intervention that has already proven potential to impact the gut-brain axis. For instance, administration of *Lactobacillus rhamnosus* GG during the first 6 months of life has been shown to significantly reduce the incidence of Asperger syndrome (AS) or attention deficit or hyperactivity disorder (ADHD) at 13 years of age^11^. Prebiotics on the other hand are non-digestible substrates that are selectively utilized by host microorganisms^12^ that could exert benefits by modulating the indigenous gut microbiota. Along with fructo-oligosaccharides and polydextrose, galacto-oligosaccharides (GOS) are the most frequently used prebiotics in infant formula^13^. GOS has been shown to potently alter the gut microbiome by stimulating *Bifidobacterium* species^14^, key health related taxa, particularly in early life^15,16^. Importantly, GOS-induced bifidogenic shifts promote the production of microbial metabolites such as SCFAs, acetylated amino acids, and GABA, all of which are increasingly recognized for their roles in neurodevelopment. Other classes of nutrients in infant formula comprise vitamins, amino acids and minerals^17^. B-vitamins (such as B1, B2, and B6)^17^ act as essential cofactors in neurotransmitter synthesis, while amino acids like tryptophan, tyrosine, leucine, lysine, and histidine are precursors to neuroactive compounds such as serotonin, melatonin, dopamine, GABA, acetylagmatine, and pipecolinic acid. These nutrients may directly or indirectly modulate critical neurodevelopmental processes including synapse formation, myelination, and neuronal signalling. Recent findings have demonstrated that consumption of a combination of vitamins B1, B2, B6, zinc, copper, iron, histidine, isoleucine, lysine, and leucine is positively associated with increased myelination in regions of the brain implicated in social behavior^18^. Notably, prior studies indicate that approximately 10% of dietary protein—the primary source of amino acids—escapes digestion in the small intestine and is subject to fermentation by colonic microbiota^19^. Nevertheless, compared to pro- and prebiotics, the effects of these nutrients on the early-life gut microbiome remain poorly understood.

This study aimed to assess how GOS, nutrient blends (of vitamins, minerals and amino acids) and combinations thereof impact the gut microbiome of infants (2-4 months old) and young children (2-3 years old). The selection of specific micronutrients and amino acids included in the nutrient blends was based on previous work by our group, which showed that the intake of a combination of vitamins B1, B2, B6, zinc, copper, iron, histidine, isoleucine, lysine and leucine was positively correlated with levels of myelination in social brain areas in infants^18^.

This prior evidence provided a strong rationale for testing these ingredients, not only for their nutritional value but also for their potential roles in supporting early neurodevelopment via microbiome-mediated and direct mechanisms.

To obtain mechanistic insights, we used the *ex vivo* SIFR^®^ technology, a high-throughput, miniaturized, bioreactor-based model that accurately cultivates the gut microbiome in the laboratory. Importantly, this technology has been validated to provide clinically predictive insights in microbiological^20^ and host endpoints^21^. Our findings underscore the prebiotic potential of GOS in early life, while highlighting how nutrient blends can further potentialize prebiotic effects on a broad spectrum of microbiome-derived neuroactive metabolites.

## Materials and methods

### Test products

GOS (GOS-950-P) was provided by King Probiotics (Guangdong, China). Nutrient blends (BL for infants, BL+ for young children) were provided by Nestlé Research (Lausanne, Switzerland). The composition and test doses of the nutrient blends are provided in **Table 1**. GOS was tested at 4 g/L.

**Table 1.** Nutritional composition (g/L) of the blends of vitamins, micronutrients and amino acids for each age group.

| Ingredient | BL (Infants) | BL+ (Young children) |
| --- | --- | --- |
| Vitamin B1 | 0.00115 | 0.00027 |
| Vitamin B2 | 0.00175 | 0.00040 |
| Vitamin B6 | 0.00158 | 0.00036 |
| Zinc | 0.00936 | 0.00215 |
| Iron | 0.01458 | 0.00334 |
| Copper | 0.00095 | 0.00022 |
| Histidine | 0.36728 | 0.08429 |
| Isoleucine | 0.88298 | 0.20265 |
| Lysine | 0.88261 | 0.20257 |
| Leucine | 1.59484 | 0.36603 |
| **Total** | **3.75708** | **0.86228** |

### Fecal donor selection criteria

Fecal samples were collected according to a procedure approved by the Ethics Committee of the University Hospital Ghent (reference number BC-09977; approval date 13/4/2021). Parents of infants and young children signed an informed consent in which they donated the fecal sample of their child for the current study.

Inclusion criteria for 2-4 months-old infants were exclusive formula-feeding (without human milk oligosaccharides), while 2-3 years-old young children had to comply to an omnivorous diet. Exclusion criteria were antibiotic use in the past 3 months and gastrointestinal disorders (cancer, ulcers, IBD). This resulted in the enrolment of 12 specific subjects (n = 6 per age group) with an average age of 3.0 (± 0.6) and 29.5 (± 3.3) months for infants and young children, respectively.

### Ex vivo intestinal fermentation assay (SIFR^®^) design

Four study arms were evaluated for each age group, i.e. GOS, nutrient blends (BL (infants), BL+ (young children)) the combination thereof (GB (infants), GB+ (young children)) and an unsupplemented parallel control with a minimal growth medium (NSC, no substrate control) (**Figure 1A**). This NSC served as reference for evaluating treatment effects as it is identical to the treatments (same microbiome, same nutritional medium), except for the presence of GOS and/or blends.

**Figure 1.**
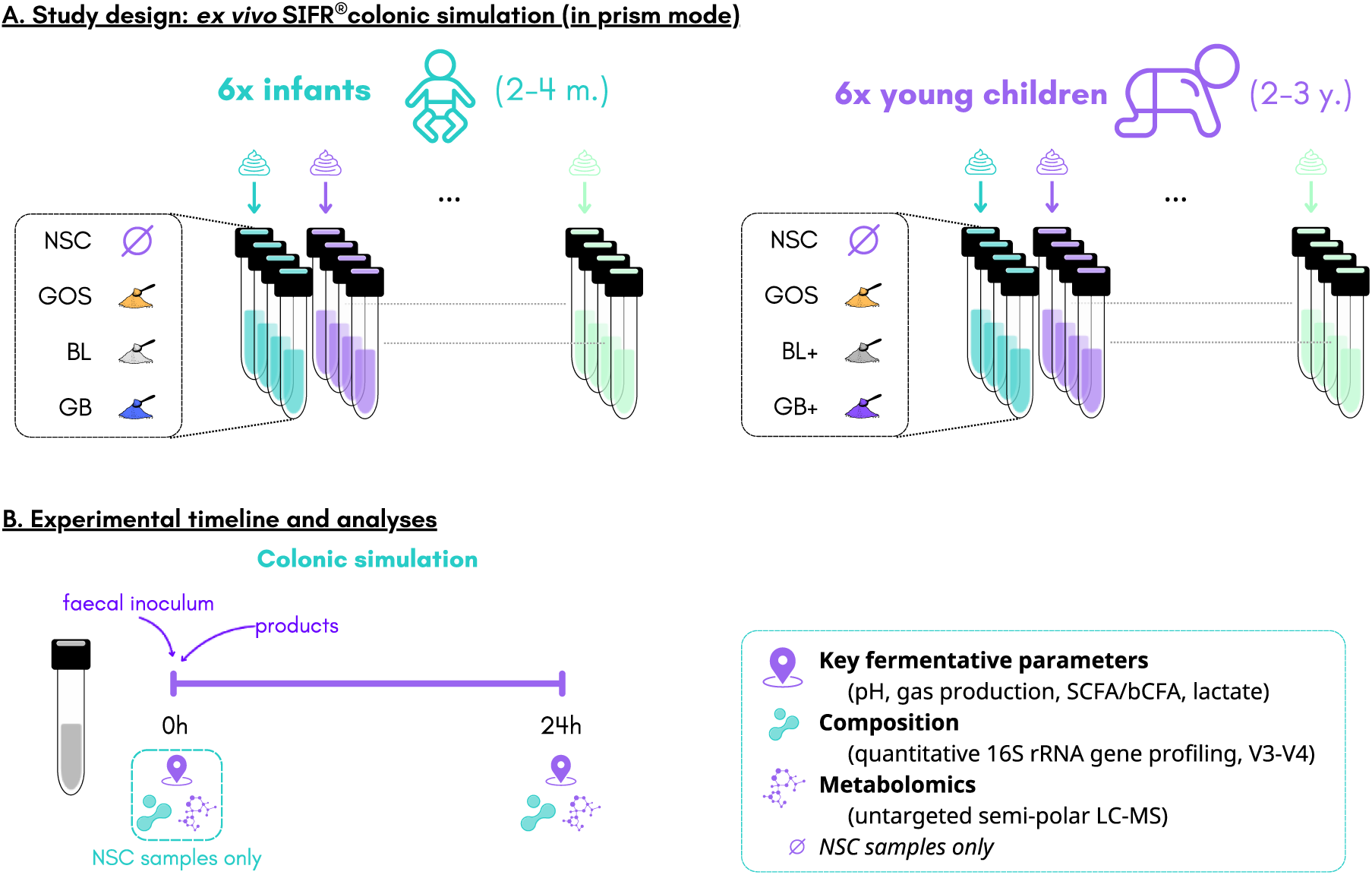
The impact of GOS, nutrient blends (BL (infants), BL+ (young children)) and combinations thereof (GB (infants), GB+ (young children)) were assessed on the gut microbiome of infants and young children using the *ex vivo* SIFR^®^ technology. (a) Study design along with (b) timeline and analysis.

Colonic fermentation was studied using a standardized *ex vivo* fermentation system with the SIFR^®^ technology as previously described^20^. Briefly, individual stool samples were processed in a bioreactor management device (Cryptobiotix, Ghent, Belgium). Each bioreactor contained 5 mL of a mixture of nutritional medium (M0019 (infants) or M0024 (young children), Cryptobiotix, Ghent, Belgium), fecal inoculum and test product. After being rendered anaerobic, bioreactors were incubated under continuous agitation (140 rpm) at 37°C. Upon gas pressure measurement, samples were collected at 0 h and 24 h for measurement of key fermentative parameters (pH, gas, SCFA, BCFA and lactate production), microbial composition and in-depth metabolite analysis (**Figure 1B)**.

Control samples (NSC) were run in technical triplicate for each infant and child with analysis of key fermentative parameters confirming coefficients of variation below 3% illustrating the high technical reproducibility of the SIFR^®^ technology.

### Key fermentative parameters

As previously described^22^, after extraction in diethyl ether, SCFA (acetate, propionate, butyrate) and branched-chain fatty acids (BCFA; sum of isobutyrate, isovalerate and isocaproate) were determined via gas chromatography with flame ionization detection. Lactate was quantified using an enzymatic method (Enzytec^TM^, R-Biopharm, Darmstadt, Germany). Finally, pH was measured using an electrode (Hannah Instruments Edge HI2002, Temse, Belgium).

### Taxonomic microbiota analysis by quantitative 16S rRNA gene profiling

Quantitative insights were obtained by correcting proportions (%; 16S rRNA gene profiling) with total counts (cells/mL; flow cytometry), resulting in estimated cells/mL of different phyla, families and OTUs (operational taxonomic units), as recently described^22^. Briefly, DNA was extracted using the SPINeasy DNA Kit for Soil (MP Biomedicals, Eschwege, Germany), according to manufacturer’s instructions. Subsequently, library preparation and sequencing were performed on an Illumina MiSeq platform with v3 chemistry. 16S rRNA gene V3-V4 hypervariable regions were amplified using primers 341F and 785Rmod. To determine total bacterial cell density, samples were diluted and subsequently stained with SYTO 16 after which cells were counted via a BD FACS Verse flow cytometer (BD, Erembodegem, Belgium).

### Untargeted metabolite profiling

As described in a recent publication^23^, liquid chromatography–mass spectrometry (LC-MS) analysis was carried out on a Thermo Scientific Vanquish LC coupled to Thermo Q Exactive HF MS (Thermo Scientific), using an electrospray ionization source, both in negative and positive ionization mode. The data analysis focused on level 1 and 2a metabolites given the high degree of certainty of the annotation: level 1 (annotation based on retention times (compared against in-house authentic standards), accurate mass (with an accepted deviation of 3ppm), and MS/MS spectra), level 2a (annotation based on retention times and accurate mass). An additional criterium to focus on microbiome-derived metabolites was to restrict the main statistical analysis on those metabolites that exhibited intensity (expressed as area under curve, AUC) from 0 h (NSC) to 24 h (any study arm) for at least for 4 of the 6 test subjects (within a given age group). This approach ensures that the analysis targets metabolites actively produced by the gut microbiota, rather than those that were consumed between 0 h and 24 h.

### Data analysis

All analyses were performed using R (version 4.2.2; www.r-project.org; accessed on 4 November 2024). R software was used to make principal component analysis (PCA), violin plots and heat maps. For the key fermentative parameters and metabolomics analysis, significance of treatment effects was assessed via repeated measures ANOVA analyses (based on paired testing among the six subjects per age group), with p-value correction according to Benjamini-Hochberg^24^. For the statistical analysis of microbial composition, three measures were taken. First, the analysis was performed on log_2_-transformed values. Second, a value of a given taxonomic group below the limit of detection (LOD) was considered equal to the LOD, as previously described^20^. Finally, paired t-tests were performed on the 50 and 100 most abundant OTUs which covered on average 99.8% and 98.8% of the abundances of a given sample for infants and young children, respectively. To unravel microbial taxa responsible for the production of specific metabolites, Regularized Canonical Correlation Analysis (rCCA) was executed using the mixOmics package with the shrinkage method for estimation of penalization parameters (version 6.20.3)^25^.

## Results

### Microbiome composition was age-related

First evidence of age-dependent differences in fecal microbiome composition was that α-diversity (Chao1 diversity index) was considerably lower for infants (33.0 ± 11.2), compared to young children (128.1 ± 19.7). Further, interpersonal differences in fecal microbiome composition were noted amongst infants (Figure 2A), with *Bifidobacteriaceae* levels being as abundant as 75% of the community for infant 4 while being absent for infant 3, whose microbiome consisted of *Enterobacteriaceae* and *Clostridiaceae* (Figure 2B). In contrast, interpersonal differences among young children were smaller with dominant community members belonging to the *Bacteroidaceae, Lachnospiraceae* and *Ruminococcaceae* families. The *Bifidobacteriaceae* family was consistently present for young children at levels from 2.2 to 6.2%.

**Figure 2.**
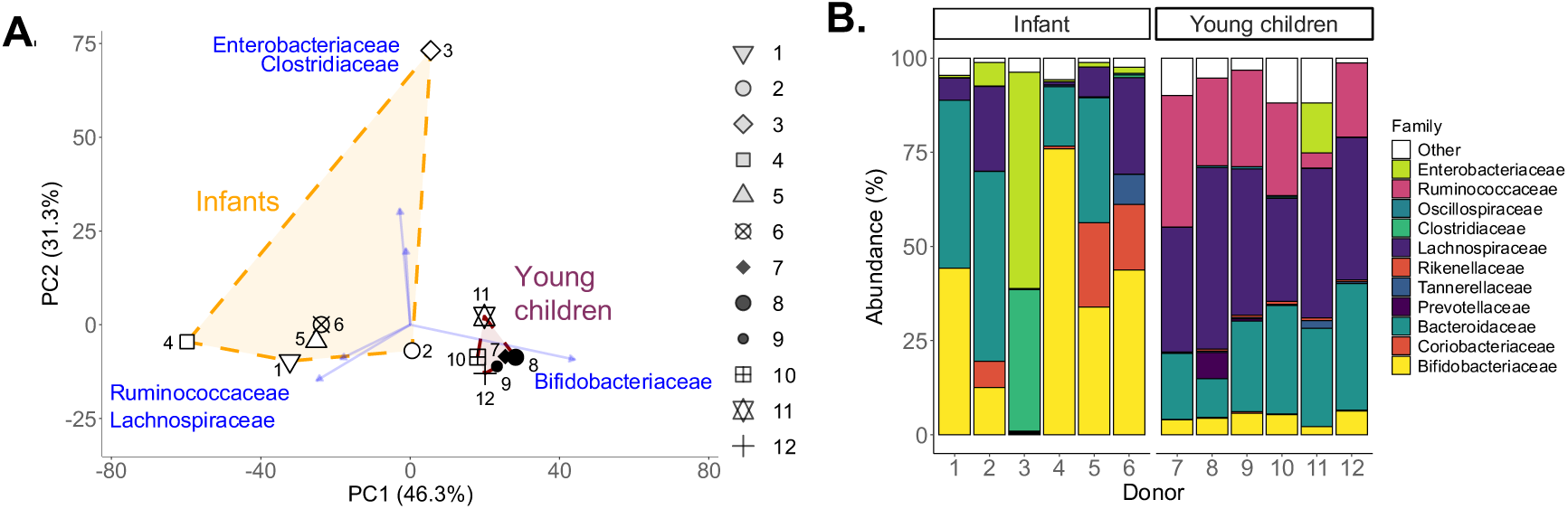
Microbiome composition was highly age-dependent, with profound interpersonal differences among infants, contrasting with the adult-like microbiota of young children. (A) PCA based on centered log_2_-transformed abundances of families (%), highlighting the 5 families that explained most variation in the dataset. (B) Abundances (%) of most abundant families.

### GOS-containing treatments stimulated SCFA, while nutrient blends rather enhanced BCFA

GOS was well fermented by the microbiota of infants and young children as followed from the stimulated acetate levels for both age groups (especially infants) (Figure 3A,G). While GOS also enhanced propionate for both age groups (Figure 3C,I), it increased lactate specifically for infants (Figure 3B) and butyrate specifically for young children (Figure 3J).

**Figure 3.**
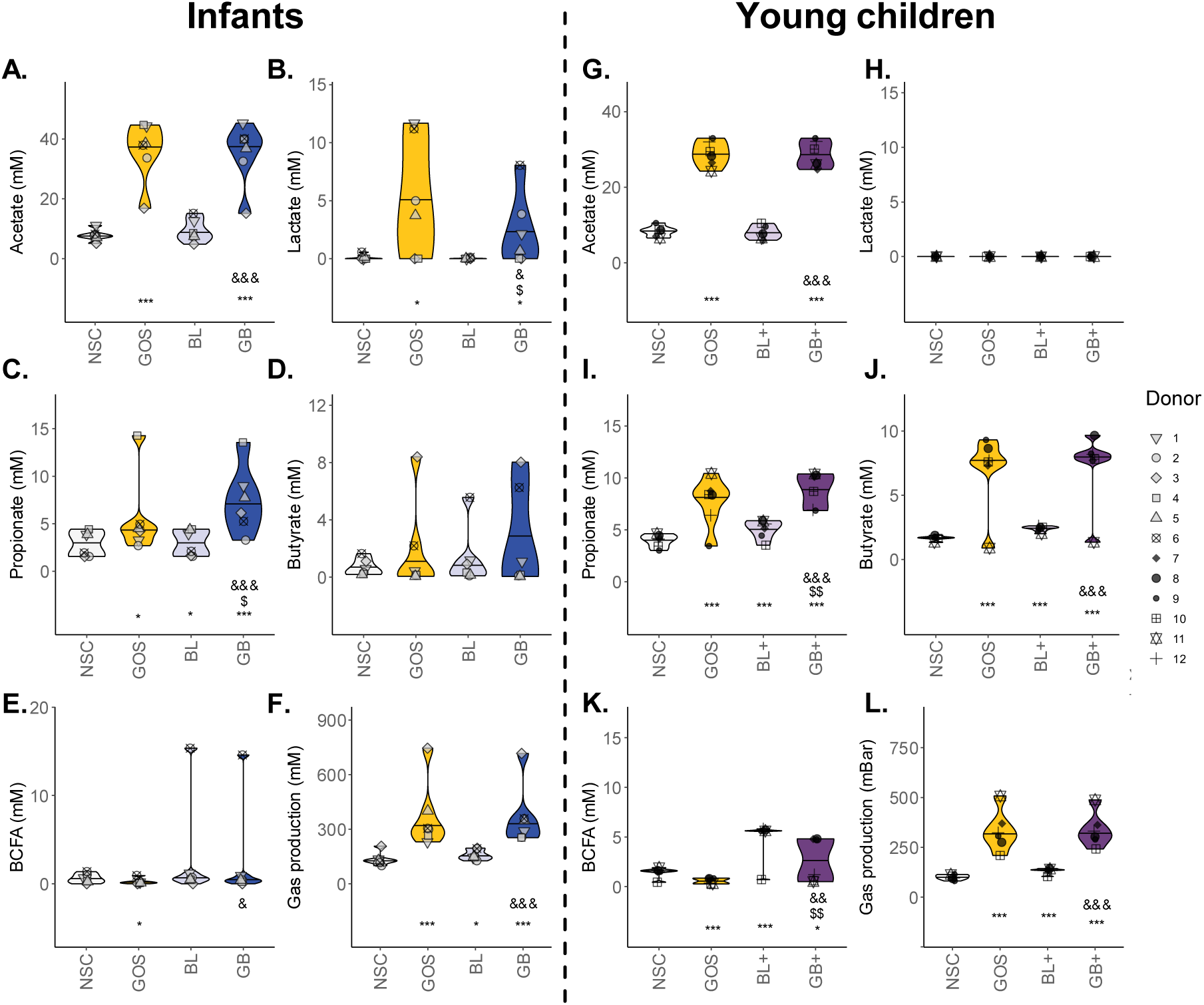
GOS-containing treatments (GOS, GB, GB+) stimulated SCFA, while nutrient blends (BL, BL+) rather enhanced BCFA (mainly for young children). SCFA/BCFA/lactate (mM) and gas production (mbar) for (A-F) infants and (G-L) young children upon treatment with nutrient blends (BL, BL+) and combinations thereof (GB, GB+) compared to an unsupplemented control (NSC). Statistical differences with NSC are indicated with * (0.10 < p_adjusted_ < 0.20), ** (0.05 < p_adjusted_ < 0.10) or *** (p_adjusted_ < 0.05). ‘$’ indicate differences between GOS and GB/GB+, while ‘&’ indicate differences between BL/BL+ and GB/GB+.

Additionally, the nutrient blend BL+ significantly stimulated propionate, butyrate and especially BCFA, while BL only mildly but significantly increased propionate production for the infants (Figure 3).

Combinations of GOS and BL or BL+ (GB or GB+) resulted in highest propionate and butyrate levels. While effects were mostly additive, the impact of GB on propionate for infants revealed a synergy between GOS and BL as the impact of GB exceeded the sum of the individual effects of GOS and BL (Figure 3C).

Finally, two remarkable interpersonal differences in butyrate production were noted for infants (Figure 3D): GOS or GB strongly increased butyrate for infant 3, while butyrate along with BCFA were especially notable with BL or GB for infant 6 (Figure 3E).

### GOS and nutrient blend altered gut microbiome composition of infants and young children in a product-specific manner

GOS, both in absence and presence of nutrient blends (GOS, GB, GB+), impacted the microbiota of both age groups by boosting *Bifidobacteriaceae* (Figure 4A,B). This followed from the stimulation of OTUs related to a broad range of species including *B. longum*, *B. bifidum* and especially *B. pseudocatenulatum* (Figure 4C,D) that correlated to elevated acetate levels upon GOS treatment (Figure 5A,B).

**Figure 4.**
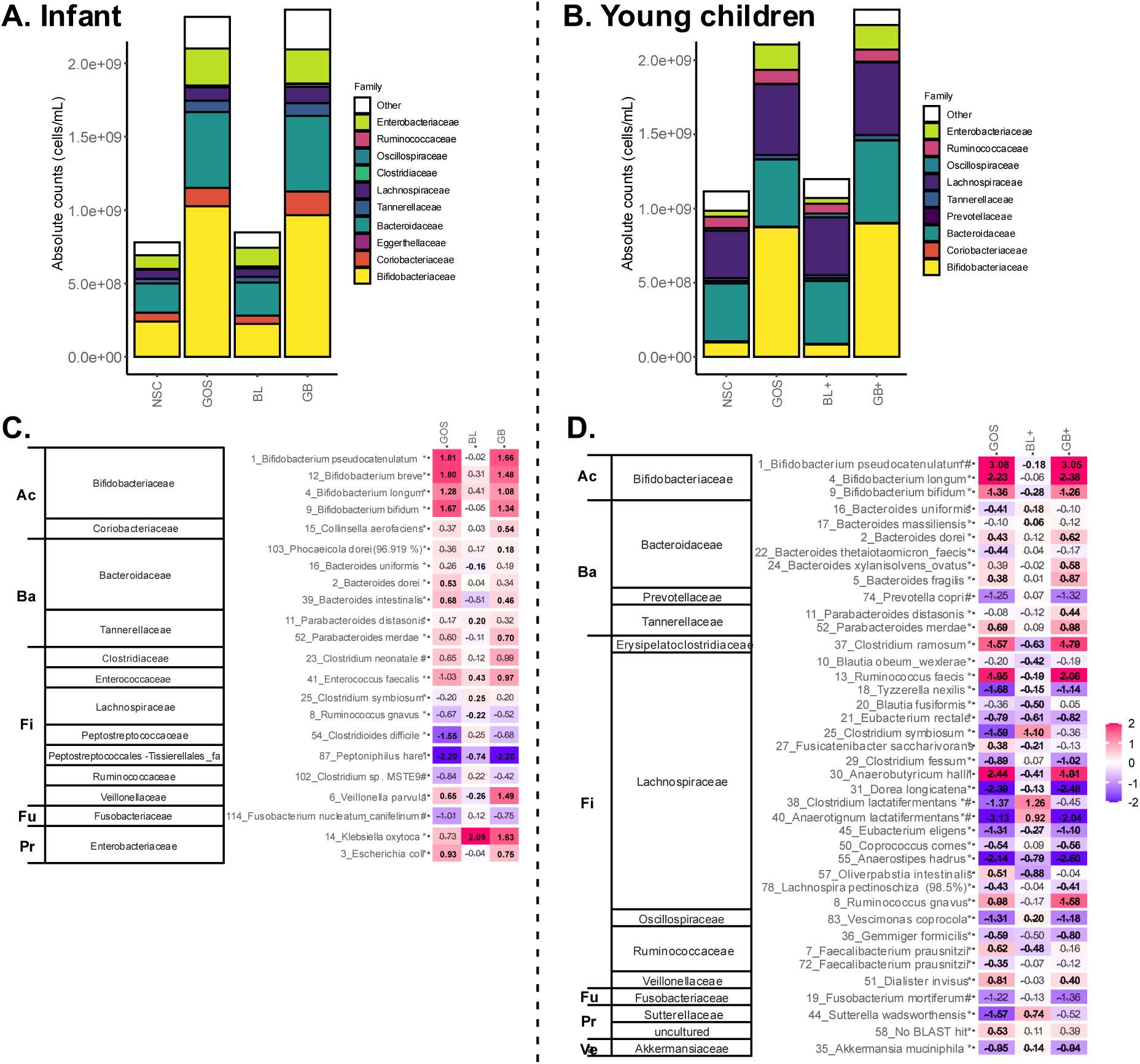
GOS strongly impacted gut microbiome composition, strongly enhancing *Bifidobacteriaceae*, both in absence (GOS) and presence of nutrient blends (GB, GB+). (A) Absolute levels (cells/mL) of the most abundant families for infants and (B) young children upon treatment with nutrient blends (BL, BL+) and combinations thereof (GB, GB+) compared to an unsupplemented control (NSC). Heatmap for (C) infants and (D) young children based on OTUs that were significantly (*p* < 0.20) affected by any of the treatments (highlighted with ‘*’) or that explained most variation in a PCA analysis (highlighted with ‘#”; PCA not shown), expressed as log_2_ (treatment/NSC), averaged over all test subjects within an age group. Values indicated in bold show significant increases (> 0) or decreases (< 0). Phyla to which the families belong are indicated on the left (Ac = Actinobacteriota, Ba = Bacteroidota, Fi = Firmicutes, Fu = Fusobacteriota, Pr = Proteobacteria, Ve = Verrucomicrobiota).

**Figure 5.**
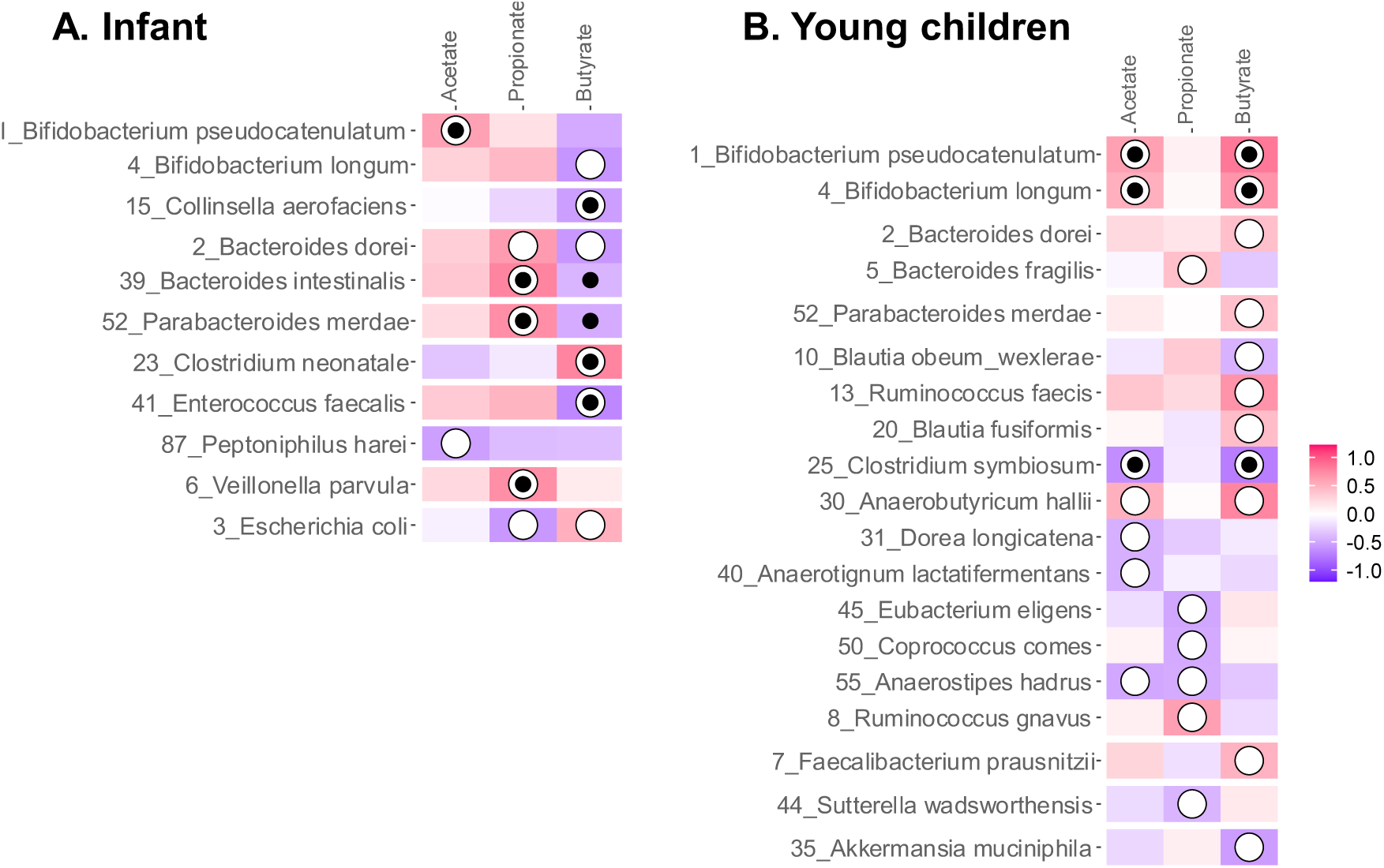
Regularized Canonical Correlation Analysis (rCCA) to highlight correlations between SCFA or BCFA and microbial composition across all study arms for (A) infants (threshold = 0.46) and (B) young children (threshold = 0.41). The white circles indicate values larger than the threshold and black dots indicate statistical significance (*p* < 0.05) in the individual correlations between the SCFA and the OTUs based on Spearman’s rank correlation coefficient.

For infants, GOS-containing products also promoted *Bacteroidaceae, Tannerellaceae* and *Veillonellaceae* (Figure 4A) due to increases of OTUs related to *Bacteroides dorei*, *Bacteroides intestinalis*, *Parabacteroides merdae* and *Veillonella parvula* (Figure 4C) that correlated with elevated propionate levels (Figure 5A). GOS or GB boosted butyrate for one infant (infant 3) related to potent increases of *Clostridium neonatale* for this specific infant. Besides stimulating various taxa, GOS-containing treatments lowered the abundance of, amongst others, an OTU related to pathogenic *Clostridioides difficile*.

For young children, GOS and GB+ stimulated OTU related to *Ruminococcus gnavus* which was associated with elevated propionate production and *Bifidobacterium pseudocatenulatum, Ruminococcus faecis* and *Anaerobutyricum hallii* – all three correlating with increased butyrate levels (Figure 4D, Figure 5B). Besides, *Faecalibacterium prausnitzii* (OTU7) which was promoted by GOS was also positively correlated with butyrate in the rCCA analysis (Figure 4D, Figure 5B).

For both donor groups, GOS also increased *Enterobacteriaceae* members, an effect that was mainly driven by increases for test subjects where GOS did not exert bifidogenic effects, i.e. infant 3 and child 11 that had no or low *Bifidobacteriaceae* levels at baseline (Figure 2).

BL or BL+ exerted milder effects. Strongest effects were noted for young children for which BL+ increased the abundance of OTUs related to *Anaerotignum lactatifermentans* and *Clostridium lactatifermentans* (Figure 4D) that related to elevated BCFA levels (Figure 5B). Additionally, in infants’ microbiota, BL significantly increased *Clostridium symbiosum* (OTU25) (Figure 4C) which positively correlated to butyrate based on Spearman’s rank correlation coefficient analysis (Figure 5A).

### GOS and nutrient blends altered the gut metabolome of infants and young children

An in-depth metabolomic analysis confirmed marked effects on metabolite production (Figure 6). In both age groups, GOS exerted the strongest impact and promoted health-related metabolites derived from amino acid fermentation including 2-hydroxyisocaproic acid (HICA), N-acetylated amino acids, aromatic lactic acids (3-phenyllactic acid, hydroxyphenyllactic acid, indole-3-lactic acid) as well as N-acetylspermidine and indole-3-propionic acid (young children only), alongside two additional neuroactive compounds. i.e., acetylagmatine and GABA (young children only) (Figure 7).

**Figure 6.**
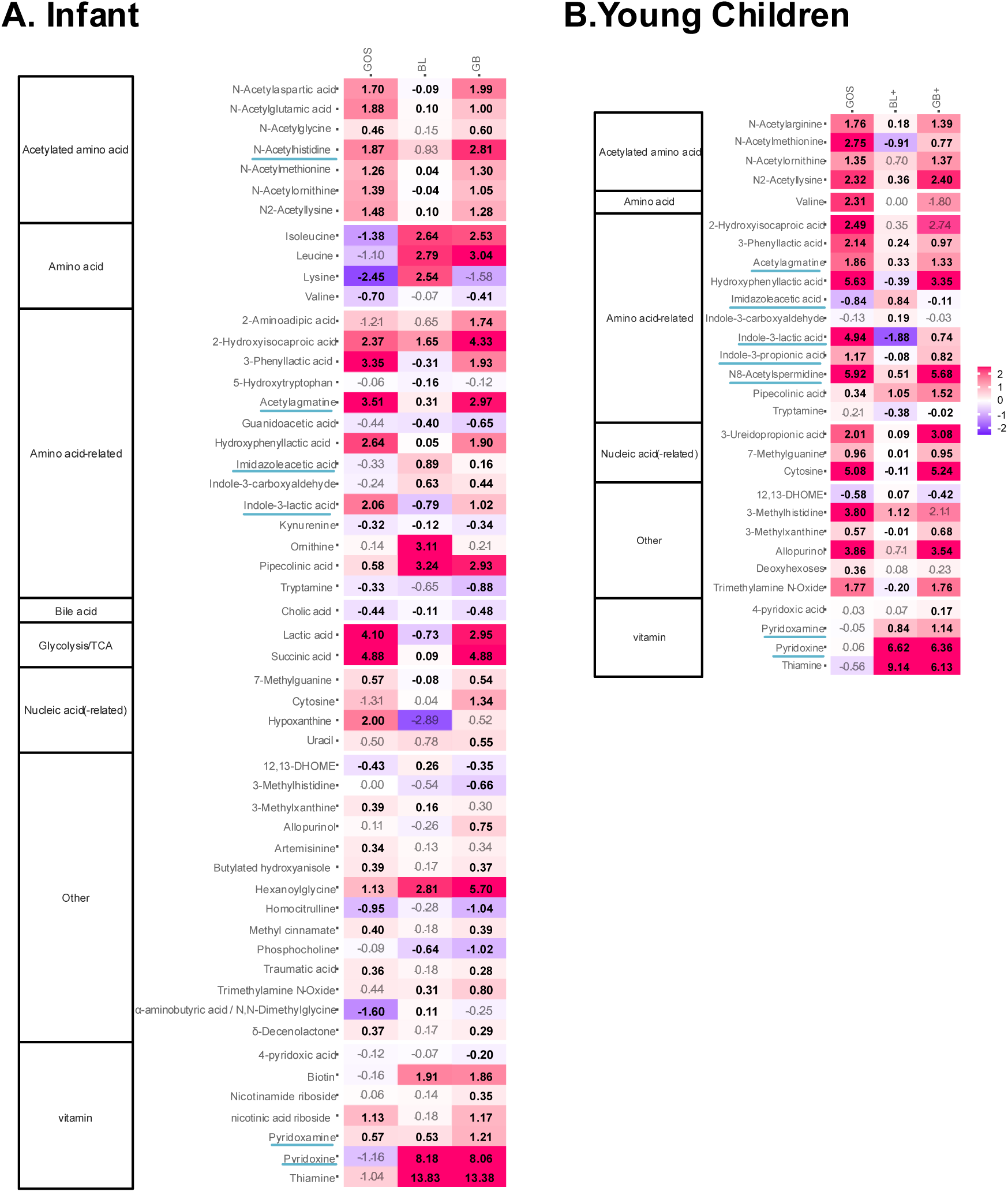
GOS, nutrient blends (BL, BL+) and combinations thereof (GB, GB+) strongly impacted the gut metabolome. Heatmap based on level 1or 2a-annotated metabolites (detected via untargeted metabolite profiling) that were significantly (*p* < 0.20) affected by any of the treatments, expressed as log_2_ (treatment/NSC), averaged over all test subjects within an age group, i.e. (A) infants and (B) young children. Values indicated in bold show significant increases (> 0) or decreases (< 0). Neuroactive or neuroprotective metabolites discussed in the text are underlined.

**Figure 7.**
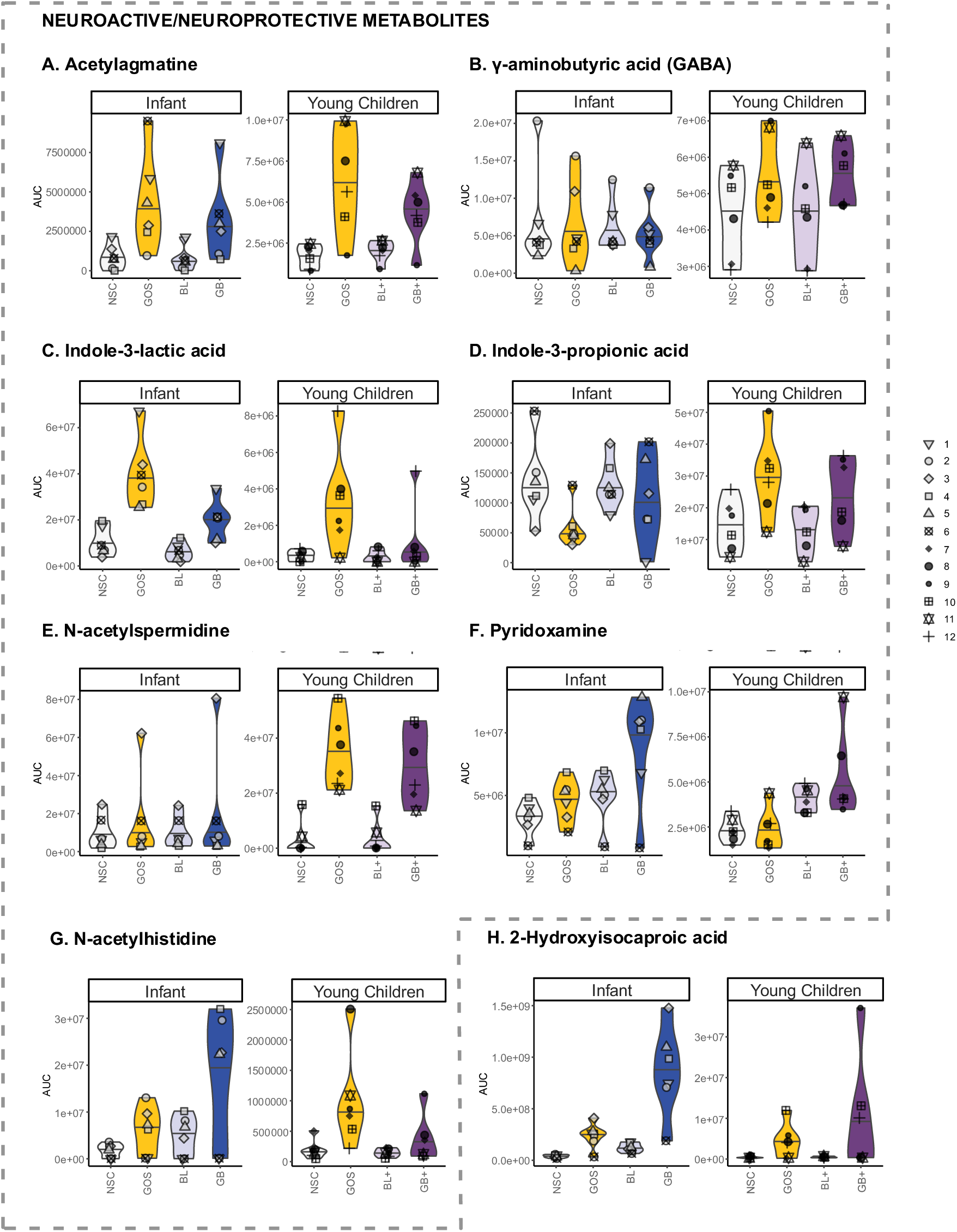
GOS stimulated neuroactive and neuroprotective compounds (A) Acetylagmatine, (B) GABA, (C) indole-3-lactic acid, (D) indole-3-propionic acid, € N-acetylspermidine, (F) pyridoxamine. Synergistic effects of combinations of GOS with nutrient blends (GB, GB+) were noted on (F) pyridoxamine, (G) N-acetylhistidine and (H) 2-hydroxyisocaproic acid. Absolute levels of these metabolites (AUC = area under the curve), as detected via untargeted metabolite profiling.

Further, for both age groups, BL and BL+ increased metabolites such as indole-3-carboxaldehyde^26^, imidazoleacetic acid^27,28^, pipecolinic acid^29,30^ (Figure 6). In addition, levels of different B-vitamins, i.e., biotin (infants only), pyridoxine (vit B6) and thiamine (vit B1), were also elevated (Figure 6). Strikingly, synergistic effects between nutrient blends and GOS were noted for HICA (GB or GB+), N-acetylhistidine (GB) and pyridoxamine (GB) (Figure 7F, G, H).

## Discussion

### Summary of main findings in perspective to literature

First, the infants and young children displayed clear age-related differences in fecal microbiome composition, in line with previous studies: while the infant microbiome was predominantly colonized with *Bacteroidaceae, Enterobacteriaceae* and particularly *Bifidobacteriaceae,* young children had a more adult-like microbiota characterized by a high diversity of microbes mostly belonging to the *Bacteroidaceae, Lachnospiraceae* and *Ruminococcaceae*^7,8^. Moreover, strong interpersonal differences, particularly among infants, enabled obtaining representative insights into how GOS, blends and combinations thereof impact the microbiome of infants and young children.

Age-related and interpersonal differences impacted the outcome of the treatments. First, the stimulation of propionate, butyrate and BCFA production by the nutrient blends was only consistent (and significant) for young children, demonstrating that its more mature microbiota allows for better fermentation of the nutrient blends. Key constituents of the blends were lysine, that can indeed be converted to propionate^31^ and butyrate^32^, and branched chain amino acids (leucine, isoleucine) that are known to be metabolised to isovaleric acid^32^ (main BCFA detected during the current study). A key species stimulated by nutrient blends for young children was *Anaerotignum lactatifermentans* (formerly *Clostridium lactatifermentans*) that has indeed been shown to ferment leucine or isoleucine to BCFA^33^. This highlights how age-related differences in microbiome composition affect treatment outcomes, stressing the importance of developing age-tailored nutritional interventions.

Another age-related finding was that propionate was synergistically promoted by the combination of GOS with the nutrient blend for infants. As reviewed by Silva *et al.* (2020)^2^, SCFA like propionate impact gut-brain communication via multiple pathways, e.g. (i) influencing immunity and gut barrier integrity upon binding to G protein-coupled receptors, (ii) interacting with receptors on enteroendocrine cells to induce the secretion of gut hormones thereby promoting signalling to the brain, or (iii) affecting other tissues upon entering the systemic circulation. The latter includes crossing the blood-brain barrier, impacting the integrity thereof and for instance contributing to the biosynthesis of serotonin^2^. Additionally, higher fecal propionate levels were previously associated with longer uninterrupted infant sleep^34^. Another example of synergistic effects between GOS and nutrient blends was the strong stimulation of HICA in both age groups. Mechanistically, HICA is a metabolite of leucine, indicating that GOS shifts microbial fermentation from isovalerate (a BCFA) towards HICA. HICA is primarily produced by lactic acid bacteria like the bifidobacteria that were increased by GOS. Additionally, HICA was shown to have antimicrobial^35^ and anti-inflammatory activity^36^, while it can also improve muscle recovery^37^. Additional synergistic effects on N-acetylhistidine (infants) and pyridoxamine (young children) further highlight the potential of the combination of GOS with nutrient blends to maximize metabolic activity of the gut microbiome. Pyridoxamine is a form of vitamin B6 which is critical in neurotransmitter synthesis (e.g., GABA, serotonin, dopamine)^38^, suggesting that its microbial enhancement could influence early neural signalling pathways.

Further, for both age groups, BL increased indole-3-carboxaldehyde (tryptophan-derived metabolite linked to improved gut barrier function^39^), imidazoleacetic acid (histidine-derived neuroregulatory metabolite^28^), pipecolinic acid (lysine-derived linked to beneficial effects such as alleviating constipation^40,41^) along with levels of different B-vitamins (biotin, for infants only), pyridoxine and thiamine, highlighting the added value of providing nutrient blends in early life.

Despite age-related differences in microbiome composition, GOS consistently promoted *Bifidobacteriaceae* for both age groups, and boosted the production of health-related SCFA, in line with earlier findings in children in similar age range^14,42^. This demonstrates rather predictable outcomes, even if the starting microbiome composition is highly different. This bifidogenic effect related to the stimulation of a broad range of (potential neuroactive) metabolites including acetate, the key SCFA produced by *Bifidobacterium* species^43^. Besides SCFA, GOS stimulated acetylagmatine (arginine metabolite), the acetylated form of agmatine that has neuroprotective effects against acute but also chronic neurodegenerative diseases^44^. In an animal model, agmatine was also implicated in mood regulation and synaptic plasticity, making it a compelling metabolite of interest in early-life interventions^45^. The acetylated form of spermidine (N-acetylspermidine) is also stimulated by GOS for the young children. Spermidine, a polyamine derived from agmatine, has been shown to have protective effects against neuroinflammation and neural damage in different models^46–48^. Further, for young children, GOS also stimulated GABA production, a glutamate-derived, inhibitory neurotransmitter with relaxing, anti-anxiety, and anti-convulsive effects^49^. GOS also stimulated tryptophan, tyrosine and phenylalanine-derived aromatic lactic acids which is produced by infant-type *Bifidobacterium* species associated with breastfeeding and have been linked to immune health and gut barrier function^15^. Especially, indole-3-propionic acid, a derivative of indole-3-lactic acid promoted by GOS, can readily enter the circulation and cross the blood-brain barrier, and has been intensively studied for its neuroprotective effects^50^. In animal models, it promotes neurogenesis and recovery after axonal injury^51^, while in humans, its levels are positively associated with brain-derived neurotrophic factor^52^ that support the growth, survival, and differentiation of both developing and mature neurons. Additionally, indole-3-propionic acid produced by early-life gut microbiome is potentially critical in preventing allergic airway inflammation in adulthood^53^. Recent studies have demonstrated an association between stimulation of *B. longum* and increased production of indole-3-propionic acid^52,54^, likely via promoting its precursor, indole-3-lactic acid. Interestingly, indole-3-lactic acid was stimulated by GOS for both age groups and has also been shown to exhibit neuroprotective effects^55^. Altogether, the stimulation of these metabolites highlights potential health benefits of GOS in pediatric nutrition.

Another notable finding was that butyrate production in infants and young children involved different taxa. For young children, butyrate production positively associated with to the presence of *Bifidobacterium pseudocatenulatum,* along with *Anaerobutyricum hallii*, a species that is unable to ferment oligosaccharides but can potently cross-feed with *Bifidobacterium* species to produce butyrate (via acetate or lactate)^56^ or propionate (via 1,2-propanediol)^57^. In contrast to the young children, *A. hallii* was absent in the infant microbiota. Instead, individual infants showed butyrate production induced by GOS potentially via stimulation of *Clostridium neonatale,* as evidenced by its positive correlation with butyrate levels (Figure 5A). *C. neonatale* has not been demonstrated to cross-feed *Bifidobacterium* spp. to enhance butyrate synthesis. Altogether, these findings contribute to the literature on succession of butyrate-producing taxa during infant gut microbiota development^58,59^.

### Strengths and limitations of the study

A notable advantage of the study approach is the utilization of the *ex vivo* SIFR® technology, which has been validated to offer clinically predictive insights into gut microbiome modulation^20^ and its subsequent effects on gut barrier integrity and immune function^21^. This technology facilitates a highly controlled research setting, allowing any observed variations (except for minor technical variations) to be attributed to treatment effects. Furthermore, the SIFR® technology enables the acquisition of insights into gut-derived metabolites, which are often challenging to analyze in clinical studies due to their rapid absorption or utilization in the body. This limitation confines many clinical studies to the analysis of fecal samples, which are excretion products from which most metabolites have been removed^61,62^. Thus, these drawbacks have been addressed in the current study. Importantly, the integration of metabolomics with microbiome profiling allowed us to identify a series of neuroactive or neuromodulatory metabolites (e.g., GABA, acetylagmatine, pyridoxamine, imidazoleacetic acid, pipecolinic acid, indole-3-propionic acid) that may be relevant for early brain development. This metabolite-level resolution is a notable strength, providing a mechanistic bridge between microbiota shifts and gut-brain axis modulation. Such mechanistic insights are particularly valuable for informing the design of future nutritional interventions targeting cognitive, emotional, or behavioral outcomes in infants and young children. Nevertheless, despite the good congruence of data obtained with the SIFR^®^ technology and clinical findings^20,21^, clinical studies are required to confirm the proposed health benefits in the host. Functional validation of neuroactive metabolites—e.g., their systemic bioavailability, CNS activity, and impact on neurodevelopmental markers—remains a necessary step to move from correlation to causation.

A critical side note on the impact of GOS is that while GOS generally exerted potent bifidogenic effects in each age group, one specific infant and young child did not display bifidogenic effects. Instead, an increase of *Enterobacteriaceae*, was noted for these subjects. This suggests that the inclusion of a probiotic *Bifidobacterium* species in the formula of such young children would be beneficial to also consolidate bifidogenic effects for such donors with depleted *Bifidobacterium* levels at baseline. This observation also highlights a limitation of generalizing prebiotic effects across all infants and suggests that personalized or combined prebiotic–probiotic strategies may be more effective in achieving desired neurodevelopmental outcomes.

### Implications for future research

GOS and nutrient blends impacted microbiome-derived metabolites related to the gut-brain axis, with synergistic effects highlighting the potential of this combination to maximize benefits. Importantly, many of the metabolites modulated in this study—such as GABA, acetylagmatine, HICA, indole-3-carboxaldehyde, and pyridoxamine—are neuroactive or neuromodulatory, with emerging or established roles in neurotransmission, synaptic plasticity, neuroprotection, and the regulation of mood and behavior. This opens the door for future research to examine how these microbiota-derived compounds influence measurable neurodevelopmental outcomes, such as language acquisition, emotional regulation, cognitive performance, or myelination patterns in early childhood. It will be of interest in future studies to unravel potential benefits of these identified metabolites, especially on neurodevelopment. This could include longitudinal clinical trials incorporating neurocognitive testing, EEG or neuroimaging biomarkers, and behavioral assessments in infants or young children receiving GOS and micronutrient-enriched formulas. Furthermore, given the variability in microbiota composition and bifidogenic response among individuals, stratifying future studies based on baseline microbiome profiles may help tailor interventions to maximize neuromodulatory efficacy.

Moreover, such future studies could focus on further improving the composition of nutrient blends by including additional precursors of neuroactive compounds. The strong production of HICA from leucine in combination with GOS is a clear example of how synergies can be obtained, suggesting that this could also be valid for other gut-derived metabolites. Such additional substrates might further potentiate the effects of combinations of GOS with nutrient blends.

## Author Contributions

All authors met the criteria for authorship, and all approved this final manuscript. LF conceived the study, LF, PVDA, AK, LB and AB designed it, while PVDA supervised the conduct of the study. JP and LDV performed the data analyses and PVDA drafted this manuscript with support of LDV. All authors provided feedback and revisions on the manuscript.

## Data availability statement

The datasets generated during and/or analysed during the current study are available from the corresponding author upon reasonable request.

## Funding

This study was funded by Nestlé Research (Lausanne, Switzerland).

## Disclosure statement

LF, SKD, AK and LB are current or former employees of Société des Produits Nestlé SA (Vevey, Switzerland). AK is a current employee of DSM-Firmenisch (Kaiserhaugst, Switzterland). While the authors participated in the design of the study, the interpretation of the data, and the revision of the manuscript, they did not participate in the collection and analysis of data. PVDA, AB, JP and LDV are employees of Cryptobiotix SA.

## Notes on contributors

**Laurent Ferrier, PhD,** is a senior scientist at Nestlé Research in Lausanne, Switzerland. His area covers the physiology of the gastrointestinal tract, with a particular emphasis on the role of microbiota metabolites in gut-brain communication pathways and disorders of gut-brain interaction. He is the author of 45 scientific papers

**Shaillay Kumar Dogra, MS**, is a Microbiome Specialist at Nestlé Research in Lausanne, Switzerland. He has extensively studied early life microbiomes in diverse cohorts and interventional Clinical Trials across geographies. His focus is on understanding microbiome maturation in infants and young children, and its link with nutrition and health. He has co-authored 25 scientific papers.

**Lam Dai Vu, PhD,** is the Lead Scientist of Cryptobiotix, where he specializes in providing predictive preclinical insights into the interplay between the gut microbiome, its metabolites, and host health. With a broad scientific background, he has authored 38 scientific publications.

**Alexandros K. Kanellopoulos, PhD**, is a Neuroscientist specializing in the role of nutrition in brain development and health, working in Nestlé Research in Lausanne during the conduct of this study. With over 20 years of experience, he has authored more than 20 publications in high-impact journals such as Cell, Nature Communications, and Neuron, and has led preclinical and clinical studies on brain health in early life and aging.

**Jonas Poppe, PhD,** is a Gut Microbiome Scientist at Cryptobiotix. His research focus is on exploring strategies to steer the colonic microbial ecosystem with therapeutic outcomes, studying the gut microbiome both in vivo and in vitro. He has co-authored 11 scientific papers.

**Laurence Biehl, Food Engineer,** is a Senior Research and Development Specialist at Nestlé Product Technology Center in Konolfingen, Switzerland, with over 20 years of experience in the food industry. Her expertise encompasses food formulation, product innovation, and sensory analysis, with a strong focus on developing healthier food options that meet consumer needs

**Aurélien Baudot, MSc**, is the Chief Executive Officer at Cryptobiotix. Aside of general administration, his focus is on the translation of the research questions into technical solutions. He has extensive technical knowledge of laboratory automation, analytics, and strategic leadership.

**Pieter Van den Abbeele, PhD**, is the Chief Scientific Officer of Cryptobiotix. His scientific focus is to develop models that accurately simulate the human gut in the laboratory, thus enabling studies to decipher the impact of interventions on the gut microbiome and host physiology. He is the author of more than 100 scientific papers.

## Suppelemtentary Information

<u>Supplementary File 1.</u> Cell density measured by flow cytometry and the absolute abundances (cells/mL) of bacterial families and OTUs in the samples of infants’ and young children’s microbiota.

<u>Supplementary File 2.</u> Statistical analysis results for bacterial OTUs in the infants’ and young children’s microbiota.

<u>Supplementary File 3.</u> Canonical correlations calculated via Regularized Canonical Correlation Analysis for the three main SCFAs and the bacterial OTUs in Figure 5.

<u>Supplementary File 4.</u> Statistical analysis results for metabolites identified via untargeted metabolite profiling (LC-MS/MS).

## Notes

### Competing Interest Statement

Laurent Ferrier, Shaillay Kumar Dogra, Alexandros K. Kanellopoulos and Laurence Biehl are current or former employees of Societe des Produits Nestle SA (Vevey, Switzerland).

Alexandros K. Kanellopoulos is a current employee of DSM-Firmenisch (Kaiserhaugst, Switzterland). While the authors participated in the design of the study, the interpretation of the data, and the revision of the manuscript, they did not participate in the collection and analysis of data.

Pieter Van den Abbeele, Aurelien Baudot, Jonas Poppe and Lam Dai Vu are employees of Cryptobiotix SA.

